# Online decoding of covert speech based on the passive perception of speech

**DOI:** 10.1101/2022.11.13.516334

**Authors:** Jae Moon, Tom Chau

## Abstract

**Background:** Brain-computer interfaces (BCIs) can offer solutions to communicative impairments induced by conditions such as locked-in syndrome. While covert speech-based BCIs have garnered interest, a major issue facing their clinical translation is the collection of sufficient volumes of high signal-to-noise ratio (SNR) examples of covert speech signals which can typically induce fatigue in users. Fortuitously, investigations into the linkage between covert speech and speech perception have revealed spatiotemporal similarities suggestive of shared encoding mechanisms. Here, we sought to demonstrate that an electroencephalographic cross-condition machine learning model of speech perception and covert speech can successfully decode neural speech patterns during online BCI scenarios.

**Methods:** In the current study, ten participants underwent a dyadic protocol whereby participants perceived the audio of a randomly chosen word and then subsequently mentally rehearsed it. Eight words were used during the offline sessions and subsequently narrowed down to three classes for the online session (two words, rest). The modelling was achieved by estimating a functional mapping derived from speech perception and covert speech signals of the same speech token (features were extracted via a Riemannian approach).

**Results:** While most covert speech BCIs deal with binary and offline classifications, we report an average ternary and online BCI accuracy of 75.3% (60% chance-level), reaching up to 93% in select participants. Moreover, we found that perception-covert modelling effectively enhanced the SNR of covert speech signals correlatively to their high-frequency correspondences.

**Conclusions:** These findings may pave the way to efficient and more user-friendly data collection for passively training such BCIs. Future iterations of this BCI can lead to a combination of audiobooks and unsupervised learning to train a non-trivial vocabulary that can support proto-naturalistic communication.

**Significance Statement:** Covert speech brain-computer interfaces (BCIs) provide new communication channels. However, these BCIs face practical challenges in collecting large volumes of high-quality covert speech data which can both induce fatigue and degrade BCI performance. This study leverages the reported spatiotemporal correspondences between covert speech and speech perception by deriving a functional mapping between them. While multiclass and online covert speech classification has previously been challenging, this study reports an average ternary and online classification accuracy of 75.3%, reaching up to 93% for select participants. Moreover, the current modelling approach augmented the signal-to-noise ratio of covert speech signals correlatively to their gamma-band correspondences. The proposed approach may pave the way toward a more efficient and user-friendly method of training covert speech BCIs.

## Background

Many individuals with disabilities do not have a means of functional communication. To address this issue, brain-computer interfaces (BCIs) have stepped in to offer a solution (for a review, see (1)), but thus far, the vast majority of BCIs demand user performance of control tasks unrelated to communication (e.g. a popular task being motor imagery), making BCIs unintuitive and mentally taxing to use. Of the many tasks surrounding BCIs, covert speech has become increasingly popular in the context of restoring communication. Covert speech (CS) is the mental imagery of speaking without articulation (2,3). It is often referred to as inner speech, an operationalized term in neurolinguistics for language-related thoughts (4–6). This task has been linked to a diverse set of neurocognitive functions including reading, writing, planning, and memory (7–9). Due to its ubiquity, CS has been gaining popularity in the fields of neurosciences and BCI. CS is a promising task in speech restoration as it is intuitive and naturally aligned with the functional goal of communication (rather than doing a separate unrelated task to control a communication aid). The development of BCIs on the basis of CS has only recently begun to garner momentum and interest, with studies revealing promising results on classifying units of CS (10–14). For instance, studies have demonstrated the potential of decoding neural patterns relating to the imagery of consonants and vowels (14, 15), phonemes (16,17), syllables (18), whole words (19,20), and sentences (21).

Although the theoretical potential of CS BCIs is undisputed, this task faces a tremendous barrier to practical translation. Specifically, it is challenging to collect large volumes of reliable data corresponding to CS. Over an extended period of time, the task itself, i.e., repeated mental rehearsal of words without feedback, is fatiguing and not conducive to sustaining user attention or engagement. Lack of reliable training data then leads to degradation of BCI performance (22,23). It is thus currently difficult to create a CS BCI for naturalistic communication with a non-trivial vocabulary, requiring a more accessible and user-friendly training paradigm. Moreover, most work on existing CS BCIs appear to focus on offline operation (for a review, see (24)) which cannot be easily extended to real-time operation. Although many decoding strategies have been implemented in CS BCIs, there appears to be large variability in performance and success rates likely due to inconsistency and lower signal-to-noise ratio (SNR) of CS data (24,25). Due to these issues, a strictly CS paradigm offers limited BCI control.

To address the low reliability of CS training signals, we propose a CS BCI trained in part on signals pertaining to passive speech perception (SP). The hypothesis that CS can be classified on the basis of SP stems from the extensive evidence of the bidirectional linkage between production and perception systems of speech (i.e., action-perception coupling) (26–30). Although the two processes constitute counter directional processes, there has been accumulating reports of shared activations in speech processing regions of the brain where the fundamental contrastive speech units are represented (26–36). Importantly, recent studies have revealed that SP and CS exhibit temporally correlated patterns of activation in shared and distinct cohorts of cortical areas, potentially suggesting that both tasks may similarly engage core computations for processing language (33,37). Given these correlations across tasks, a cross-condition model trained to accurately classify CS based on SP appears plausible and may eventually reduce the burden on users. Further progress on this research may potentially lead to the training of CS BCIs using audiobooks which, coupled with unsupervised learning, could eventually support proto-naturalistic communication. However, there appears to be a paucity of prototypical implementations of such a SP-CS model.

Here, we present an online, ternary BCI incorporating a SP-CS cross-task learning model. Fundamentally, we hypothesized that the discrimination of covertly rehearsed words could be achieved by analyzing the correspondence of brain signals in SP and CS. In the current study, we performed ternary classification with a two-layer hierarchical architecture: CS vs. rest (task-level) and if CS, then speech item 1 vs. speech item 2 (word-level). The protocol involved a unique dyadic setup whereby in each trial of the experiment, participants heard one of eight words (‘arch’, ‘archbishop’, ’gran’, ’grandchildren’, ’kin’, ‘kindergarten’, ‘mar’, ‘marketplace’) and subsequently mentally rehearsed them upon cue. The two most distinguishable words were then chosen via offline analysis. Analytically, the signals of the CS portion of a dyad were linearly transformed via a function derived from both SP and CS data (i.e., Ridge regression). Covariance matrices were extracted from the transformed CS data, and subsequently projected onto a tangent space, forming a feature vector. The best performing classifier (among logistic regression, ridge regression, and support vector machine) and regularization strength were chosen based on a grid search with cross-validation. Classifying between two words and resting state data, the online operation of this BCI yielded up to 93% accuracy, averaging between 75-80% and exceeding the chance level of 60%. Importantly, we found that the proposed perception-covert modelling effectively enhanced the SNR of CS signals with token-based specificity (i.e., SP and CS of the same word) and seemingly dependent upon the γ-band correspondence between tasks.

## Materials and Methods

### Participants

Ten native English-speaking participants aged 20-40 years, without disabilities or known health conditions (6 females, 3 males, 1 N/A, 31.4 years old ± 5.8) were recruited for this study. All participants were right-handed to ensure consistency of hemispheric dominance with respect to language processing. Age, sex, handedness, ethnicity, previous experience with BCI studies (yes/no), and fluency in other languages were documented. Participants provided informed written consent. The study protocol was approved by the research ethics boards of Holland Bloorview Kid’s Rehabilitation Hospital and the University of Toronto.

### Instrumentation

We used a 128-channel ActiCap EEG CAP (brainproducts GmbH). 64 channels were utilized (Fig. 1) with a focus on temporal, central, and parietal regions and ground and reference electrodes at AFz and FCz, respectively. Channels Fp1 and Fp2 were used as ocular artifact detectors and were excluded from analyses. Data were sampled at 1000Hz and collected through BrainVision recorder (Brain Products GmbH) and a lab streaming layer (Schwartz Center for Computational Neuroscience, Brain Products GmbH).

**Figure 1.**
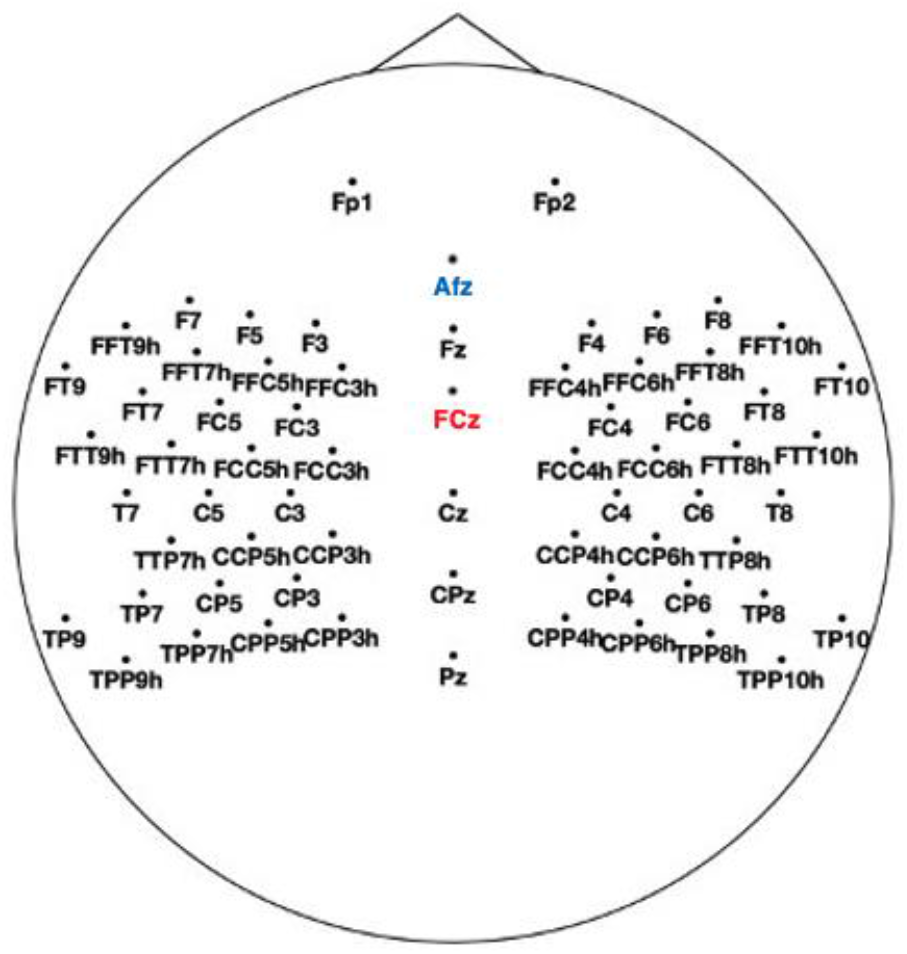
EEG montage.

Given its millisecond temporal resolution and our interest in leveraging time-evolving properties of brain oscillations, EEG was chosen as the online BCI modality. EEG has been used extensively in speech processing research to characterise the temporal dynamics of oscillations in phase entrainment (38), processing asymmetry (39), speech intelligibility (40), semantic evaluation of speech (41), categorical processing (42), and oscillatory abnormalities in schizophrenia (43,44).

### Offline Experimental Procedure

In this experiment, SP and CS trials were presented in sequential dyads, that is participants heard a randomly chosen word and subsequently mentally rehearsed the same word (Fig. **2**). First, participants observed a blank screen with a durational jitter between 1-2 seconds, after which a green cross was displayed along with an audio of 1 of 8 words (1.5seconds). Subsequently, another blank screen was displayed (1 second), after which a red cross was shown signifying the start of CS and cueing participants to rehearse the same token that was heard (1.5seeconds). Therefore, every SP trial was followed by a CS trial with the same word. Rest trials were also dyadic in that participants heard a randomly chosen word and subsequently rested. This procedure was done to simulate an implicit, no-control rest task. Participants observed a blank screen (1-2 second jitter), followed by a green cross (1 of 8 words), a blank (1 second), the rest primer (“REST”) (2 seconds), another blank (1 second), and finally a black cross (5 seconds), signifying that the participant should rest (Fig. **2**). Additional rest time provided participants a break. Rest data were epoched at 2-3.5 seconds to limit any event related potentials that may have occurred at the beginning of the black cross. Each offline block consisted of 5 SP-CS and SP-Rest trial dyads. Each session consisted of 10 blocks yielding 50 trials/class (Fig. **3**). Participants completed two offline sessions at least 2 days apart and at roughly the same time of day. The offline sessions took approximately 1 hour.

**Figure 2.**
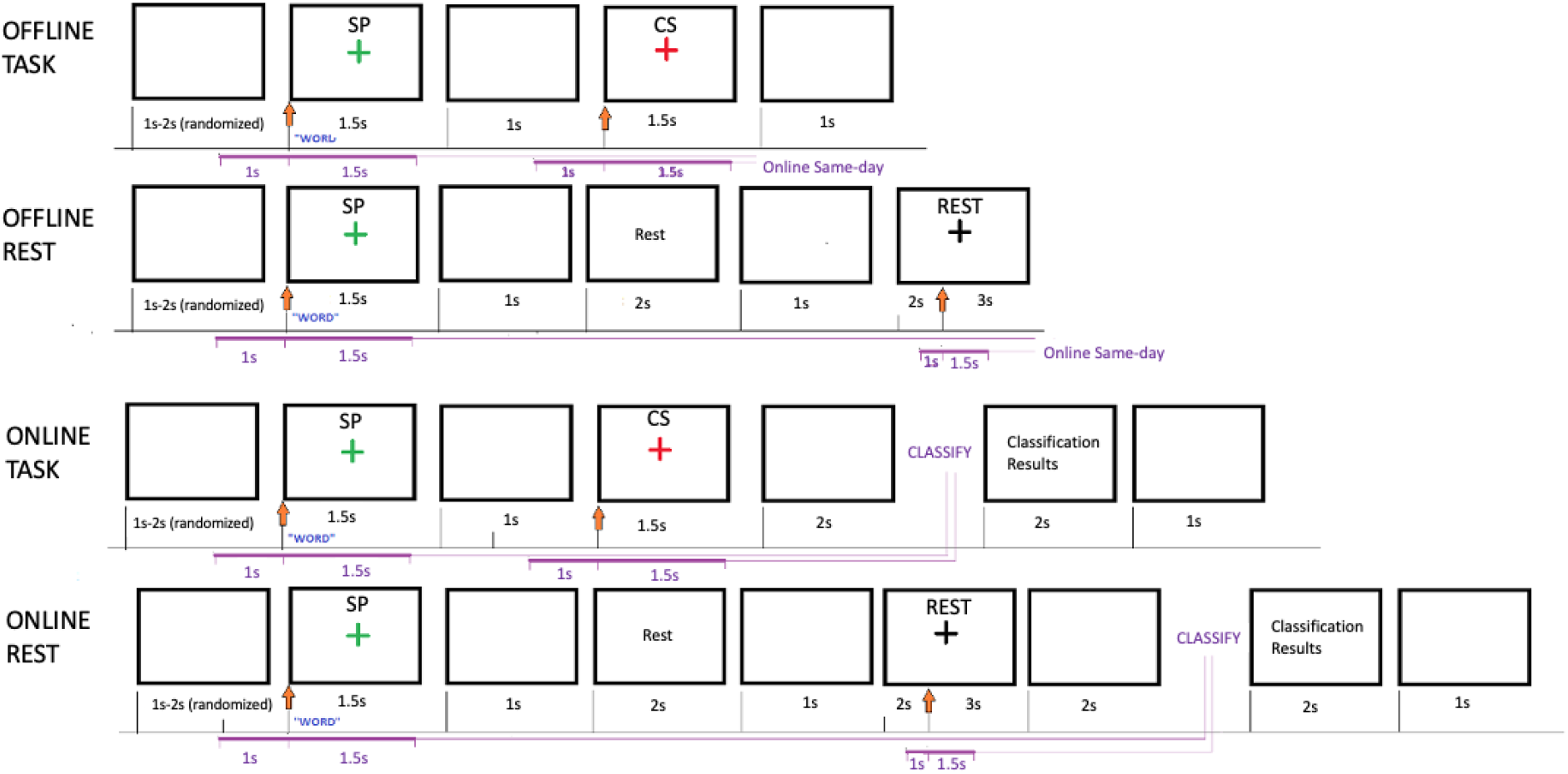
Experimental protocol. The green, red, and black cross represent SP, CS, and rest, respectively. The purple brackets depict the time range used for epoching. Blank rectangles represent denote times when participants faced a blank screen.

**Figure 3.**
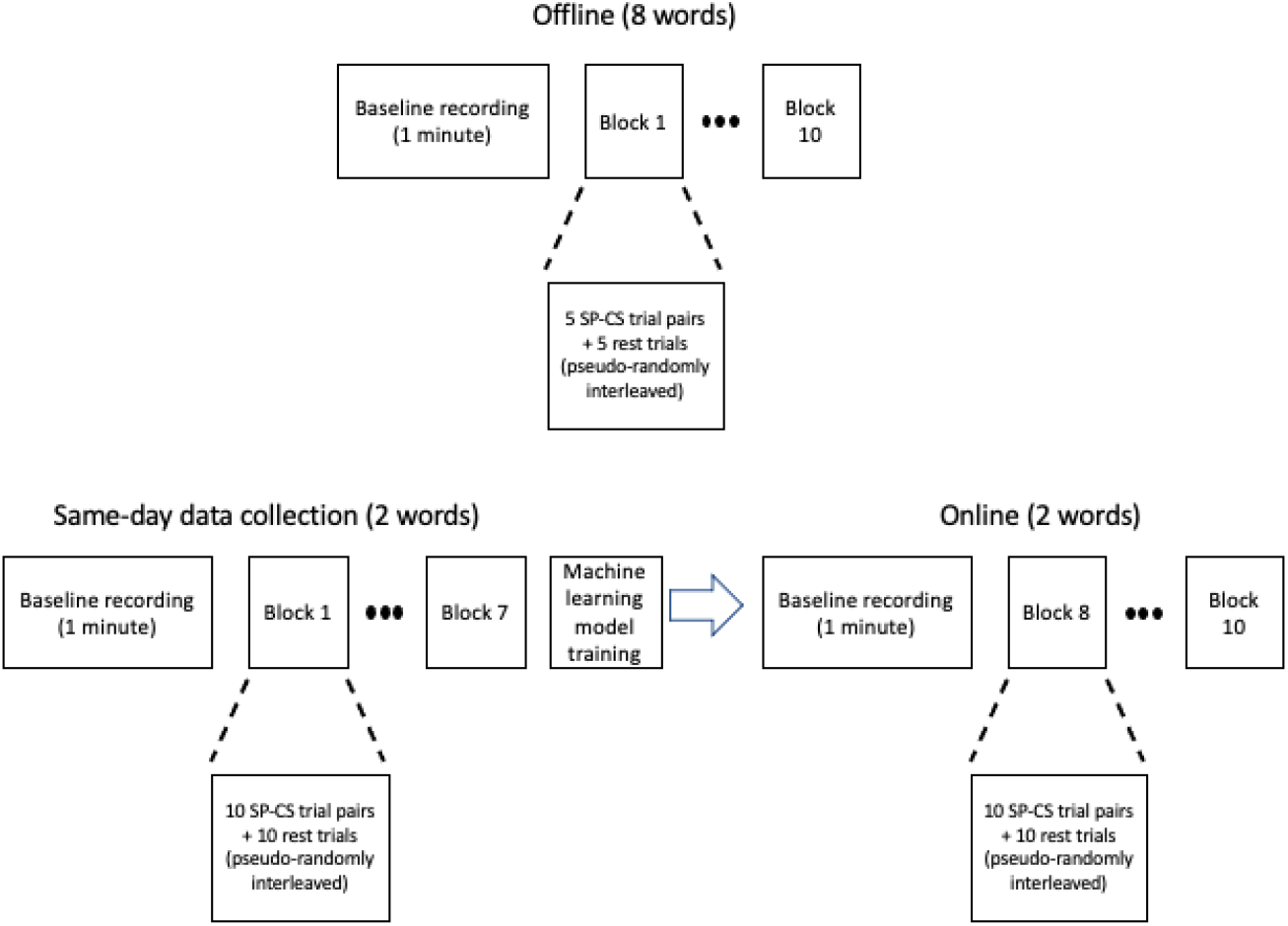
Sessional structure: offline session (top row) and online session (bottom row).

### Online Experimental Procedure

The online session (session 3) was divided into two parts: same-day data collection (offline) and online operation. A single Lab streaming layer (LSL) stream was used for both offline and online portions of session 3. The first two offline sessions were used to determine the two most distinguishable words out of the 8. The same-day and online trials exclusively used those 2 words. The protocol for same day data collection was the same as that of the previous two offline sessions with the exception that the epochs in the LSL stream were preprocessed (Fig 2). These preprocessed trials were accumulated across 7 blocks, yielding 70 trials/class. After finishing 7 blocks of offline, the screen showed a text stating: “Classifier training in progress” (Fig 3). This process took no longer than 30 seconds. After the participant-specific classifiers had been trained, participants underwent a 1-minute baseline similar to the same-day data collection protocol and the two previous offline sessions. Subsequently, the online blocks began with data being processed from the LSL stream. The online protocol (Fig. 2, bottom two rows) resembled the offline protocol but data were preprocessed and classified in real time. Therefore, after every red/black cross, participants were given classifier feedback i.e., the predicted class of the most recent trial (Fig. **2**). The classification step took no longer than 1 second and confusion matrices were updated after every trial. Each session was no longer than 1 hour.

### Speech items

There were a total of 8 words in this study: arch, archbishop, gran, grandchildren, kin, kindergarten, mar, marketplace. The 8 words were chosen from the UNION corpus^1^ containing over 16000 words. This corpus was chosen as it provides metrics such as phonology, phonological Levenshtein distance (i.e., phonological distinctness) and word frequency. For the current study, we collected monosyllabic and multisyllabic (n≥3 words) to produce better distinctions between classes.

First, the corpus was divided into one syllable and multisyllable words (n≥3 syllables). Short and long word pairs with matching phonology were selected (e.g. Arch, Archbishop), narrowing candidate words to 404 pairs. The reason for this phonology match was that the long words are filtered by high phonological distinctiveness and a matching phonology would similarly activate more specific phonological neighborhoods (45,46), encouraging more discernible signals (47,48). Audio files of words generated with google cloud to speech, approx. 150 wpm, within range of natural speech rate (49). Next, a θ oscillator-based model of syllabic segmentation was used (50) which detects boundaries using the sonority of speech. This step was necessary to ensure that audio signals had clear syllabic boundaries. A gammatone filterbank was constructed on audio signals (more in Spectrotemporal receptive field section). The oscillator’s critical damping was set to 0.5, the desired center frequency at 4Hz (51), and a threshold at 0.01. Long words with detected number of syllables equal to the amount of true syllables were selected for the next step, narrowing down the total number of words to 347 long words. Next, long words were filtered by phonological Levenshtein distance (threshold = 97.5^th^ percentile) and phonology-matching short words were filtered by frequency (lowest four among subset) to determine words that would be most neurally distinguishable. Low frequency words were preferable because they are thought to focally engage regions of interest in verbal working memory processes, as there are lesser competitors in the lexical space (52). Therefore, it was thought that neural response patterns to low frequency words would contain more robust signals due to having fewer competitors in the lexical space (53,54). With these criteria, 8 words were chosen for the experiment.

### Preprocessing

As this was an online study, preprocessing was on a single trial basis. SP and CS trials were epoched between -1 to 1.5s relative to green/red cross onset. Rest trials were initially epoched between -1 to 3.5s. After removing the mean of the 1 second baseline prior to the presentation of the black fixation cross, rest data were selected between 1s to 3.5s. For a similar approach to rest epoching see (55). (However, a post-hoc test revealed that epoching at the same time as CS relative to fixation cross onset made no significant impact on classification accuracy.) The initial sampling rate was 1000Hz. Data were filtered with a zero-phase Butterworth filter with an order of 8, low-passed at 55Hz and subsequently high-passed at 1Hz. Data were then downsampled to 256Hz. Next, we removed ocular and muscular artifacts by independent component analysis (ICA). 32 ICA components were extracted using the Picard method due to faster performance (56). Artifacts were detected and subsequently removed from the dataset based on Pearson correlation between the ICA components and filtered EEG data of selected artifactual sensors (e.g., Fp1, Fp2) (57), the thresholding (values above which a feature was classified as an outlier) of which was determined by adaptive z-scoring (57). Above threshold components were masked and the z-score was iteratively recomputed until no supra-threshold components remained. ICA weights were stored and updated cumulatively in the offline sessions and subsequently used in the online session. All preprocessing in offline, same-day data collection, and online were identical. Trial data comprised 64 channels x 640 samples (2.5s x 256Hz). The preprocessing pipeline is depicted in Figure **4**.

**Figure 4.**
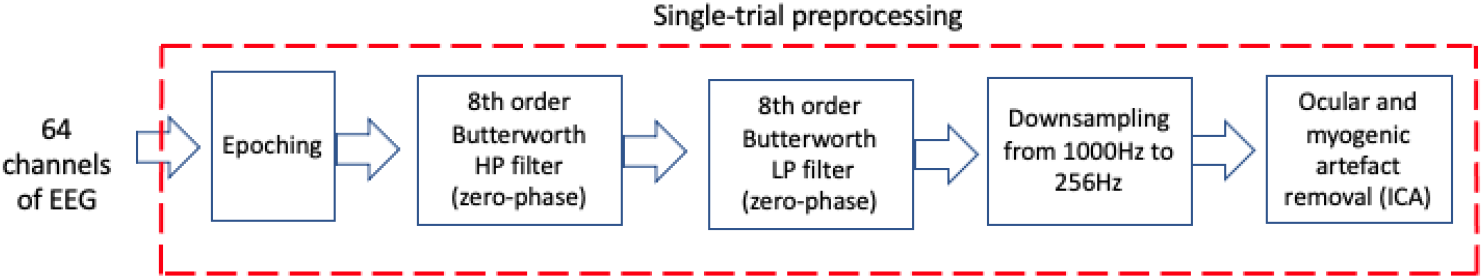
Preprocessing

### Overall classification framework

The purpose of study was to model CS based on SP such that former could be classified with brain signals from passive listening. To accomplish this, we linearly transformed the CS part of dyads using the corresponding SP trials regression. Subsequently, features were extracted using a Riemannian geometry. The two most distinguishable words were determined through an assessment of offline accuracy of each binary word pairings on a per-participant basis. These two words were used for the same-day and online portions of session 3 as well as pseudo-online analyses following sessions 1 and 2 (mimicking online situation with training-testing split of offline data). Pseudo-online and online classification were done through a hierarchical two-level architecture: 1) task-level (CS vs rest) and, if CS, 2) word-level (word 1 vs word 2). The overall classification framework is described in Figure **5**.

**Figure 5.**
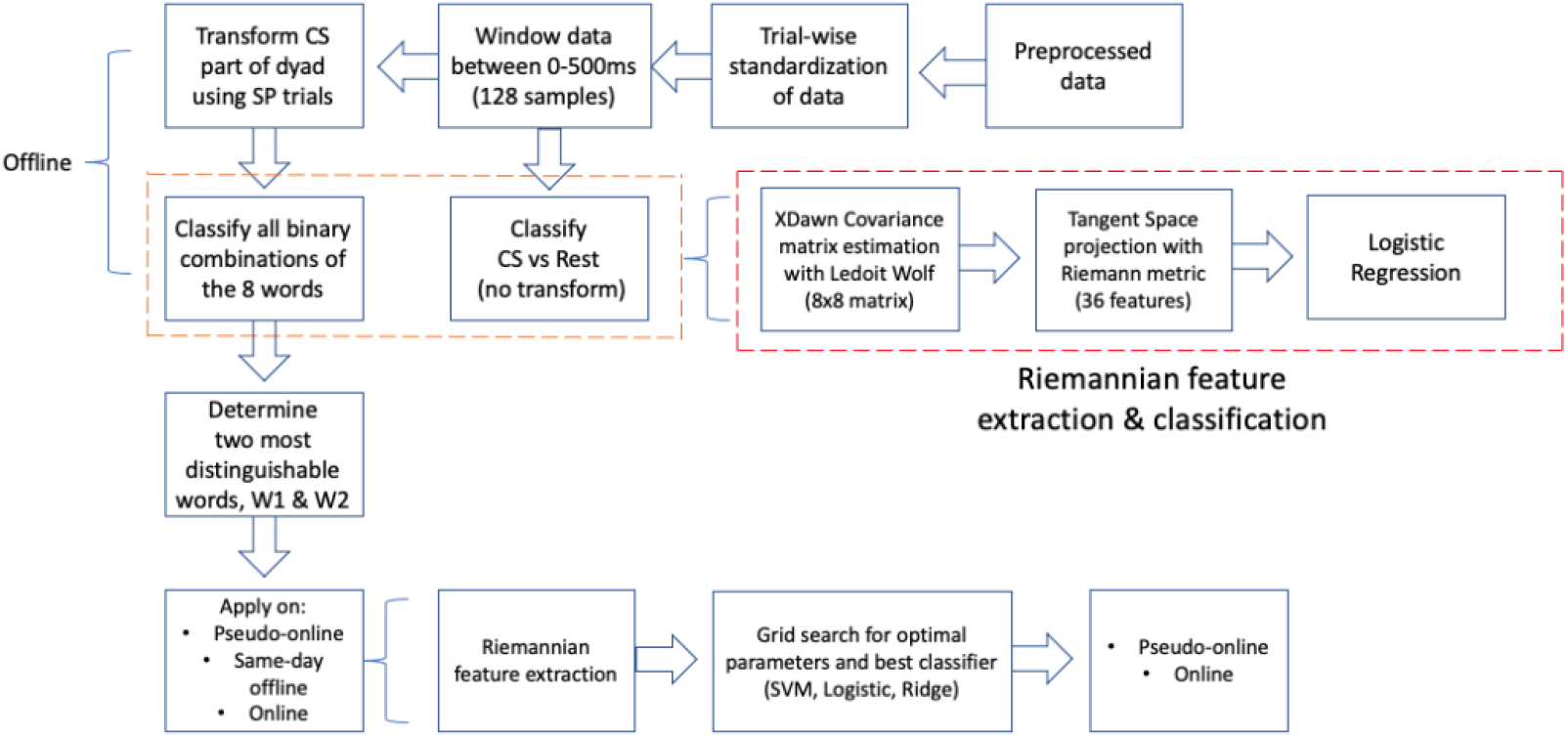
Overall classification framework.

### Transformation of CS signals

SP and CS trials were standardized (removing mean and scaling to unit variance) on a trial-by-trial basis and subsequently the first 500ms of data (128 samples) were extracted for analyses. This window of analysis empirically yielded the best accuracy averaged across folds and participants. To generate a model of CS based on SP, the CS trial of each dyad was transformed using the leading SP trial by performing ridge regression between two 64×128 matrices (target = SP, training = CS). Ridge regression was invoked because our data was highly multicollinear, which is common in EEG. With 128 samples, the ridge regression regularization parameter, alpha, was chosen based on 4-fold cross-validation (among values 0.1, 0.2, 0.5, 1, 10) on a per-trial basis (i.e., 64×128 SP and CS matrices randomly sampled into training and testing sets) and the best value was consistently found to be 0.1. This optimal alpha value determined during same-day training was applied to the online portion.

### Feature extraction

Covariance matrices were calculated on the trailing trial of dyads using the XDawn algorithm. XDawn was used because of its efficiency and accuracy with respect to extracting spatial filters of event related brain potentials and maximizing the signal to signal+noise ratio (58). The Ledoit-Wolf estimation method was invoked as the current data violated normality (Kolmogorov-Smirnov) and this method is both well-conditioned and more accurate than the typical sample covariance matrix asymptotically (59). Covariance matrices were extracted with 2 components (filters) to yield 8×8 matrices per trial. This result is obtained by the following formulation (60):

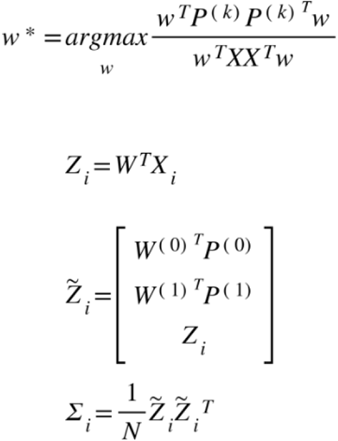

where w^*^ and W are the XDawn spatial filters derived from generalized eigenvalue decomposition combined by two class types, (0) and (1), *Z*_*i*_ is the spatially transformed data of *X*, and is a 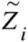 special trial structure consisting of the spatial transform of averaged evoked potentials, *P*, of classes (0) and (1), and *Z*_*i*_. *Σ*_*i*_ represents the covariance matrix with 8×8 dimensions and *N* represents the number of time samples.

Although varying numbers of components were tested, as classification typically requires 10 times as many samples as features, we opted to simplify the feature extraction with fewer spatial filters. Subsequently, the covariance matrices were projected to a tangent space whereby each matrix, belonging to a manifold, were represented in a Euclidean space (61). After projection, each matrix was represented as a vector of size *D(D+1)/2* where *D* is the dimension of the covariance matrices. Therefore, each trial yielded a feature vector of 36 elements. Feature extraction was implemented with the Pyriemann package^2^.

### Regularization and Classification

In order to determine which two words produced the most distinguishable brain responses, binary classifications between all word pairings were conducted after combining sessions 1 and 2. We used logistic regression with l-2 regularization and the optimal inverse regularization strength was determined through cross validation. For offline analyses, classification was cross validated for 4 folds with 2 repeats. The short-word and long-word pairings with the greatest overall classification accuracy were used for pseudo-online analyses, same-day data collection, and online classification.

The aggregated trials from same-day offline data collection were used to train the hierarchical online classifier. The first level of classification was between CS and Rest, prior to transformation. The second level of classification was between words. Three different classifiers were assessed: ridge classification, logistic regression, and support vector machine. The best parameters for each model type were determined via grid search. The regularization strength parameters for the models were [0.1, 0.2, 0.5, 1.0, 10.0]. Classification was cross-validated with 10 folds and the models with the greatest mean accuracy were chosen for online testing, during which the trailing trial of each dyad also underwent the same 2 classifications. The online classification workflow is depicted in Figure **6**.

**Figure 6.**
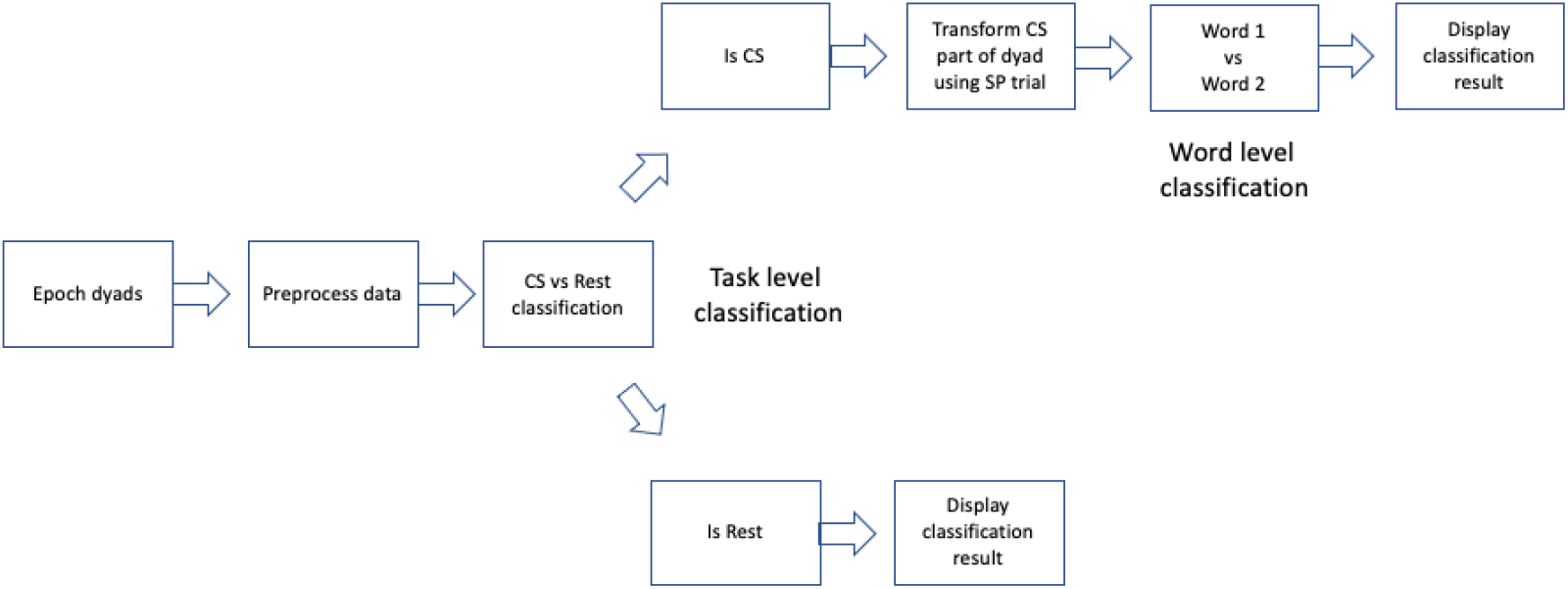
Online hierarchical classification workflow

Classification performance (accuracy and F-score) were determined for each session separately due to session-to-session variability. Accuracies across sessions were computed using Wilcoxon rank sum tests for unequal medians, as Shapiro-Wilks tests confirmed violation of normality (p<0.05). To assess the importance and specificity of the models to inherent SP-CS relationships, we repeated the above steps with 1) pure CS (no linear transformation) and 2) permuted class labels of CS. The chance level was determined by 2) to be 60%.

### Perturbation testing for determining channel contribution to classification

In order to ascertain the relative contributions of channels during classification, we performed a perturbation test retrospectively using offline SP and CS data (62). In every iteration, a given channel of the data was randomly shuffled temporally. We then measured the decrease in accuracy due to this perturbation compared to the original model with no perturbation. We repeated this procedure 1000 times to obtain a distribution of accuracy differences and subsequently computed the p-value through a z-test with alpha at 0.05, rejecting values greater or equal to this threshold. This procedure was done for every participant, and an overall topography of accuracy differences was generated by summing across participants. The greatest contributing channels were identified by highlighting the channels whose values were greater than the 99^th^ percentile of the overall distribution.

### Clustering

As the study involved the classification of 8 words in separate binary pairings, it was possible that certain words would dominate distinguishability within and among participants. To determine separability of words, we performed multidimensional scaling (MDS) and hierarchical clustering on average word event-related potentials (ERPs) of transformed CS models. Multichannel data were vectorized timewise to produce a N (8192) sample x M (8) class matrix. MDS analyses were done on these matrices of averaged word ERPs. MDS was conducted with 2 dimensions and a Euclidean metric of dissimilarity. As for hierarchical clustering, we performed Spearman Rho tests to obtain a condensed distance matrix and subsequently invoked Ward’s linkage to construct the dendrogram.

### Spectrotemporal receptive fields

In addition to hierarchical clustering and MDS, we also derived the spectrotemporal receptive fields relating to the 8 words, in an attempt to ascertain the strength of the mapping between stimulus features and neural responses. The spectrograms of the 8 words were obtained with a gammatone filterbank (63). The waveforms of the audio signals with a sampling rate of 24kHz were passed through a 64-channel gammatone auditory model filterbank (63), with lowest frequency of 50Hz and highest frequency of 12kHz. The intermediate equivalent rectangular bandwidth (ERB) filterbank matrix (64) was derived to process an audio waveform with gammatone filterbank and subsequently compute its filter coefficients. The gammatone spectrograms were then downsampled to 256Hz. Next, the average word potentials were derived by averaging across trials aggregated across participants. STRF was calculated by reverse correlation coupled with ridge regression (65):

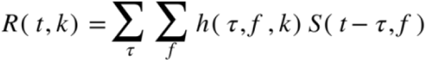

where *R*(*t, k*) is the predicted neural activity at time *t* and electrode n, 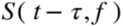 is the spectrogram representation at time *t-τ*, and acoustic frequency *f*. Finally,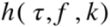 is the linear transformation matrix dependent upon the time lag *τ*, frequency *f*, and electrodes *k*. In other words, *h* represents the spectrotemporal receptive field of each electrode. Next, ridge regression with cross-validation was conducted to locate the optimal ridge parameter, α, that maximized the r^2^ score between predicted and actual neural responses. This optimal parameter value was then used to compute STRF and average across cross-validation folds. An STRF model was created for every instance of lag. To control for multiple comparisons, we conducted Monte Carlo simulation (for more information, see Section: Statistics). Topography of r^2^ values were generated from the lag at which the model maximized performance. The greatest contributing channels were identified by performing a z-test of the vectorized r^2^ scores at significance level 0.05.

### Event-related causality

To determine the relative strengths at which neural oscillations contribute to the mapping between SP and CS, we invoked a measure of causality between the two tasks known as event-related causality (ERC) as it is proficient in detecting spectral transfer in short data segments of multichannel data (66). A relationship is thought to be causal if the past information of a variable significantly improves the prediction of another variable (67). The steps to calculating ERC is can be found in (66).

ERC was calculated on preprocessed but not standardized data. First, to determine the length of time in which to calculate ERC, we conducted a series of Adfuller tests and chose the shortest sample length (186) at which data were stationary was chosen. We next determined the optimal order of the multivariate autoregressive model (MVAR) by calculating the coefficient matrix of the data and minimizing the Akaike’s information criterion (m=5). Next, we computed the cross-power spectral density from the MVAR coefficient matrix and subsequently calculated the partial coherence. The coefficient matrix was then converted to its equivalent spectral transfer function form and used in tandem with the partial coherence to derive the ERC.

ERC was calculated on all words of SP and CS observations on a per-participant basis. This yielded a 128-channel x 128-channel x 128-frequency matrix. The upper left and lower right quadrants of the ERC matrix were removed from further analyses, as these represent self-causality, and we were only interested in this measure across tasks. To determine significance level, a Monte Carlo simulation was conducted (see Section: Statistics). The total significant causality was computed for each participant for each of the 5 frequency bands by summing across channels and within frequency bands. Subsequently, the ERC of each frequency band was represented as a proportion of total ERC across all frequencies and channels. Wilcoxon rank sum tests were used to test significant differences between the causality levels in the five oscillatory frequency bands (δ, 0.2-2.5Hz; θ, 3-8Hz; α, 9-12Hz; β, 13-25Hz; γ, 30-55Hz). Topographies were generated by summing the total SP to CS and CS to SP causalities across all frequencies, or within the θ-or γ-bands. Finally, a z-test of topographical ERC values was conducted to determine significantly contributing channels (those with p<0.05).

### Event-related phase-amplitude coupling (PAC)

To understand whether the θ and γ oscillations of SP and CS dynamically adjust to ongoing speech, we determined the event-related phase-amplitude coupling (PAC) (68). For this analysis, we used preprocessed SP and CS signals grouped by long (multisyllabic) and short (monosyllabic) words. Low frequency θ phase was determined by Butterworth filtering between 3Hz to 7Hz and subsequently computing the angles of the Hilbert transform. High frequency amplitude was computed by Butterworth filtering between 30 and 60Hz with 0.25Hz widths and steps of 0.5Hz, and subsequently obtaining the absolute values of the Hilbert transform. Therefore, PAC was measured time-wise across the γ frequency range (N samples x F frequencies):

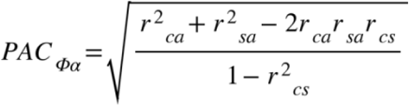

where *φ* and *a* respectively denote phase and amplitude, *r* denotes the Pearson’s correlation, and *c* and *s* denote the cosine or sine of the phase. For example, *r*_*ca*_ represents the Pearson’s correlation between the cosine of the phase and the amplitude variable.

We also examined any cross-task relationships between SP and CS by computing the degree of coupling between the θ phase of SP and the γ amplitude of CS. PAC was calculated on channel TP10, where we identified the greatest common total ERC between SP and CS. To generate the topographies, we computed PAC within every single channel and averaged across time to generate a single value per channel. PAC computation was subject to Monte Carlo simulation by randomly permutating the γ amplitude values (see Section: Statistics). To determine the sensitivity to γ amplitude patterns during PAC computation, trial-wise PAC (instead of time-wise) was computed for every trial by progressively perturbing the γ amplitude data with normally distributed noise one time sample at a time. The decay in PAC was observed with increasing amounts of contamination. The result was then fitted with an exponential decay model *a* + *be*^*−c(x)*^ (Fig. **S2**).

Finally, correlations between cross-task PAC strength and accuracy, along with a metric of γ-band dissimilarity, dynamic time-warping (DTW) (69), were tested with Spearman tests.

### Statistics

Statistical significance for STRF, ERC and ERPAC measures were determined via Monte Carlo simulations with max randomization (70). For each surrogate run, the observed data were randomly permutated, and surrogate measures were subsequently computed on the permuted data. The 95^th^ percentile value of each run was recorded. The p-values were determined as the probability at which the surrogate maxima exceeded the observed measures. 1000 runs of thisonte Carlo simulation were performed for the computation of STRF, ERC, and ERPAC as changes in p-value stabilized after this point (Δ p< 5e-4, 2e-4, 4e-4, respectively).

## Results

### Classification Results

All accuracies (Fig. 7a) and F-scores (data not shown), with the exception of offline Session 2 of Participants 5 and 6, exceeded the 60% chance level, which was determined via permuting CS class labels prior to linear transformation. This chance level estimate provided a more conservative criterion for BCI usage (where raw chance is 33%). Mean classification accuracies across all participants were above the 60% chance level, with scores not significantly different across sessions (Wilcoxon rank sum, p>0.05). Mean online accuracy was 75.3% ± 9.02. Interestingly, Session 1 accuracies were significantly greater than those of Session 2 or 7 out of 10 participants (Wilcoxon rank sum, p<0.05). Additionally, to ensure that the classification was not purely based on word duration/length, we performed binary classification between mono-or multi-syllabic words. Mean accuracy in this scenario was 53.8%. Moreover, we found no significant differences in the accuracies between participants with and without previous BCI experience (Wilcoxon rank sum, p>0.05).

**Figure 7.**
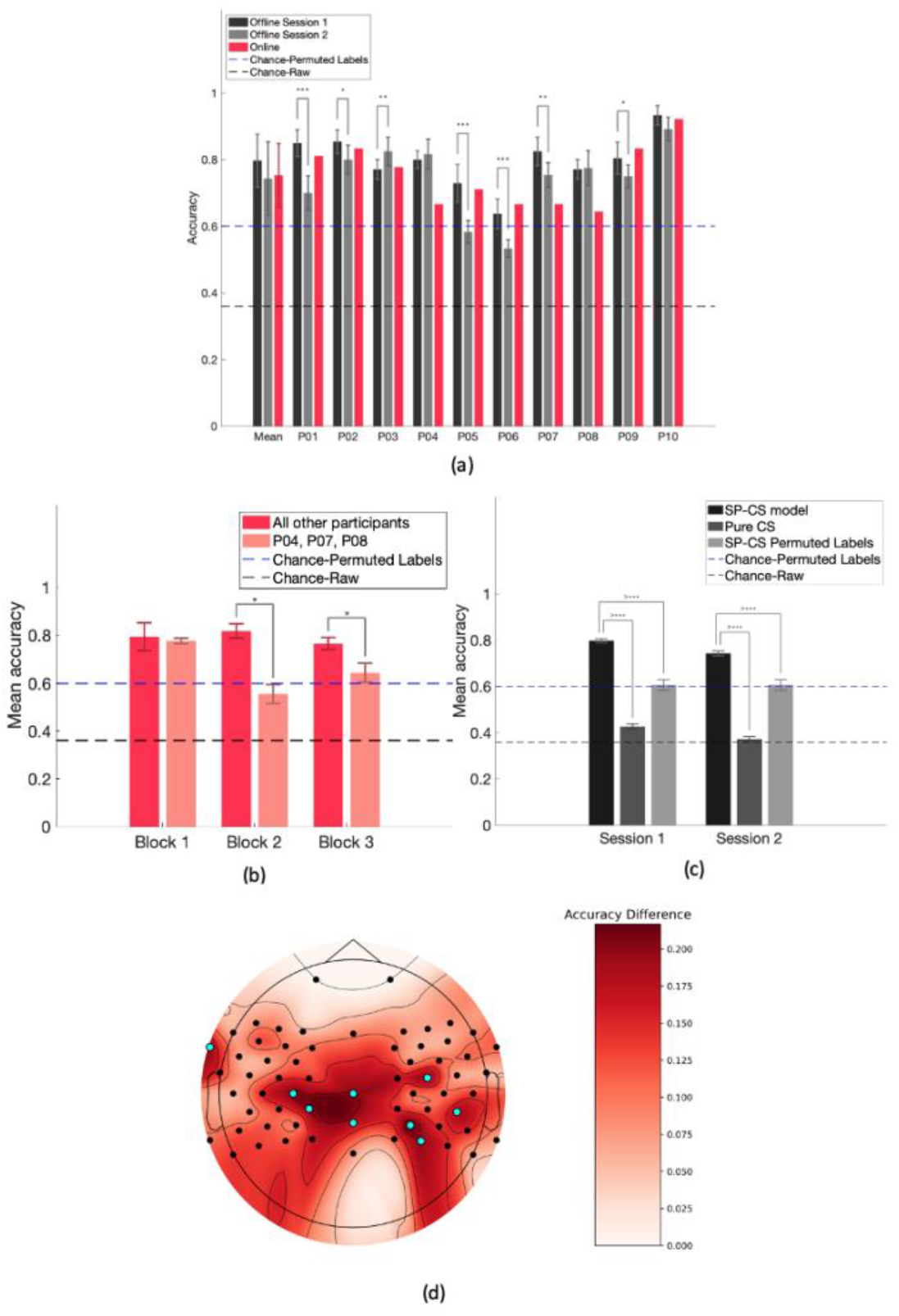
SP-CS modelling enhances classification accuracy. (a) Classification accuracies for each participant and session, including offline (black and grey bars) and online (red bars) (b) Online block-wise accuracies for participants who displayed consistent (red bars) and inconsistent (red cross-hatched bars) performance between offline and online. (c) Comparison of classifier performance between the proposed SP-CS model, a pure CS model, and one where CS transformation took place, but the trial labels were permuted, effectively disrupting the dyadic structure. (d) Perturbation test: observing the extent to which accuracy deteriorates when perturbing one channel at a time. Cyan dots depict the channels where accuracy dropped the most significantly. All significant differences were identified by Wilcoxon Rank Sum tests (***=p<0.001, **=p<0.01, *=p<0.05).

Although most participants’ online scores were consistent with their offline counterparts, Participants 4, 7, and 8 exhibited a lower online accuracy compared to their offline scores. To investigate this further, we measured block-wise online accuracy, bifurcating participants into those with consistent scores and those with inconsistent scores (Fig. 7b), with the criterion being for the latter group that the online scores were lesser than the mean-standard deviation of offline Session 1 scores. This post-hoc analysis revealed that scores were comparable between groups in block 1 but significantly deteriorated in blocks 2 and 3 for the inconsistent group (Wilcoxon rank sum, p<0.05). Overall online accuracy excluding Participants 4, 7, and 8 was 79.1% ±8.45.

Next, we compared our current modelling approach to that of a pure CS condition with no cross-condition modelling (Fig. 7c, red and black bars). This pure CS condition yielded accuracies greatly below the permutation chance level and around the raw chance level for both offline sessions. To confirm that the transformation of CS with SP did not preferentially emphasize SP-specific features during classification, we disrupted the dyadic structure of the experiment by permuting the class labels of CS prior to transformation. The accuracy arising from this permutation test was used as the theoretical chance-level. Disruption of the pairing of SP and CS of the same words led to significantly reduced classification accuracies in both sessions (Wilcoxon rank sum, p<0.0001) (Fig. 7c, red and grey bars). (A further explanation for the 60% permutation accuracy is provided in the Supplementary Text and Figure S1.)

Finally, to understand the channel-wise contribution to classification performance, we conducted a perturbation test by injecting random gaussian noise into the data one channel at a time and observing decrements in model accuracy. The highest contributors (Fig. 7d, cyan dots) to classification accuracy were the fronto-temporal, and central/centro-parietal channels.

### Separability of Words

Interestingly, ‘archbishop’ and ‘gran’ were found to be the most consistent word pairing (nine out of ten participants) associated with the highest classification accuracy, apart from Participant 6 (‘grandchildren’, ‘arch’). To understand why ‘archbishop’ and ‘gran’ were the most separable, we conducted hierarchical clustering using Ward’s linkage and multidimensional scaling (MDS) using a Euclidean distance metric (Fig. 8a,b). Both tests revealed that the neural patterns of ‘Archbishop’ and ‘Gran’ contain the largest distance between all word pairings. We next explored spectrotemporal receptive field (STRF) mapping between the gammatone spectrograms of audio signals, obtained through a filterbank of 20 logarithmically spaced frequency bands between 50Hz-12kHz according to equivalent rectangular bandwidth scale of cochlea, and neural patterns (Fig. 8c), where higher r^2^ scores depict more accurate mappings between the two. Significant r^2^ scores, determined through a z-test and depicted by the cyan markings, were observed in left frontal/posterior temporal and centro-parietal clusters, reminiscent of the perturbation test topography (Fig. 7d). Interestingly, other words exhibited a paucity of significant r^2^ scores.

**Figure 8.**
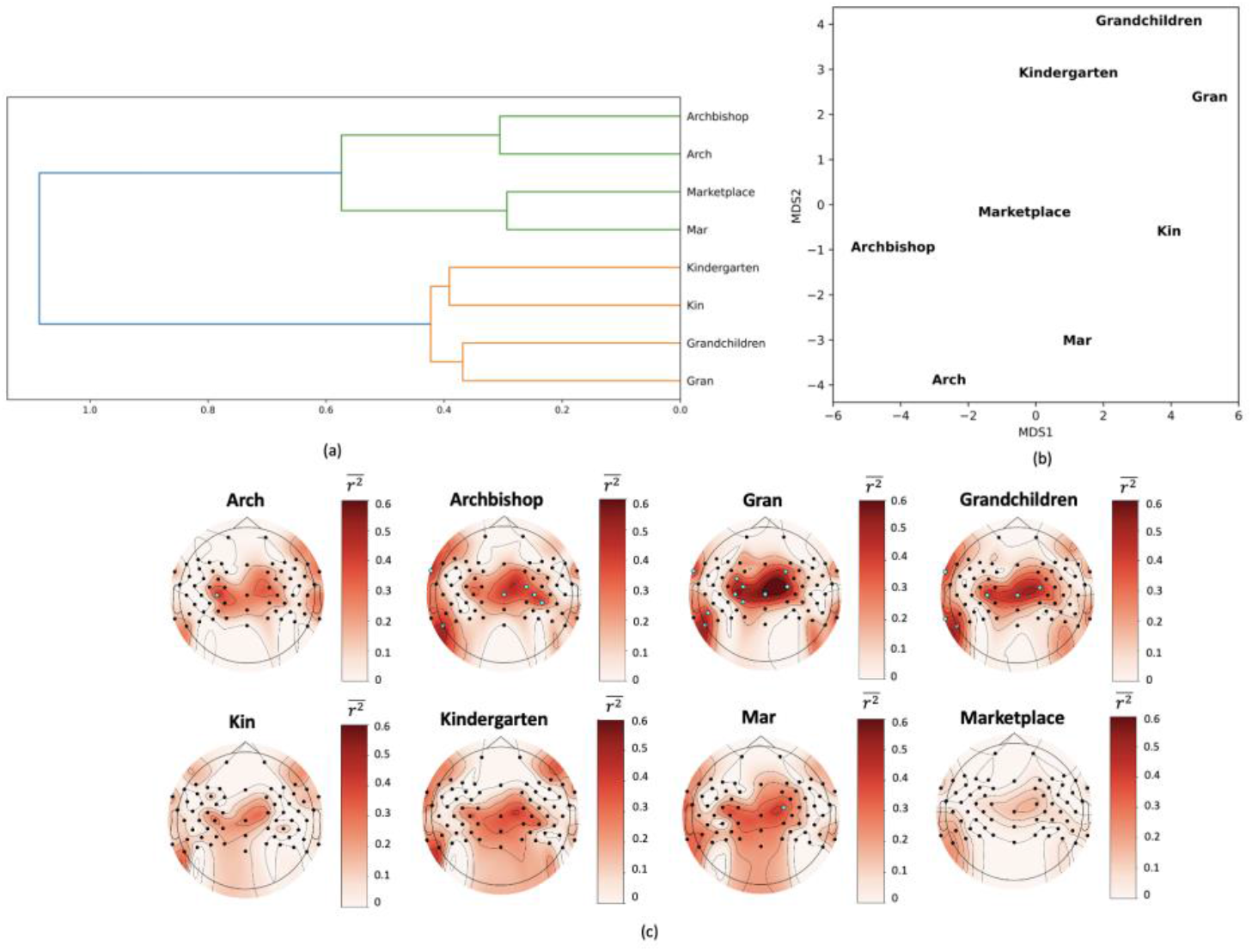
The neural patterns to ‘archbishop’ and ‘gran’ are most separable. (a) Hierarchical clustering of word-level averaged potentials including all channels using Ward’s linkage. (b) Multidimensional scaling (MDS) of word-level averaged potentials after calculating a Euclidean distance between words. (c) Spectrotemporal receptive field (STRF) topographies of r^2^ scores at best performing lags. Values denote mean r^2^ across time. Cyan markings denote significant channels through z-testing. A high r^2^ score describes a more accurate mapping between stimulus features and neural responses.

### Contributions of Neural Oscillations

Given that the modelling approach depended to some degree upon inherent relationships existing in SP and CS, we next sought to understand the relative neural oscillatory correspondence between the two tasks. Therefore, we first assessed the event-related causality (ERC), a measure of spectral causality, between SP and CS, in five frequency bands (δ, θ, α, β, γ) (Fig. 9). The majority of causality occurred in the γ-band (Fig. 9a), which was also reflected topographically (Fig. 9b,c), though evidently weaker in the CS to SP direction. Interestingly, the loci of source causality between the two tasks primarily occurred in bilateral temporal channels.

**Figure 9.**
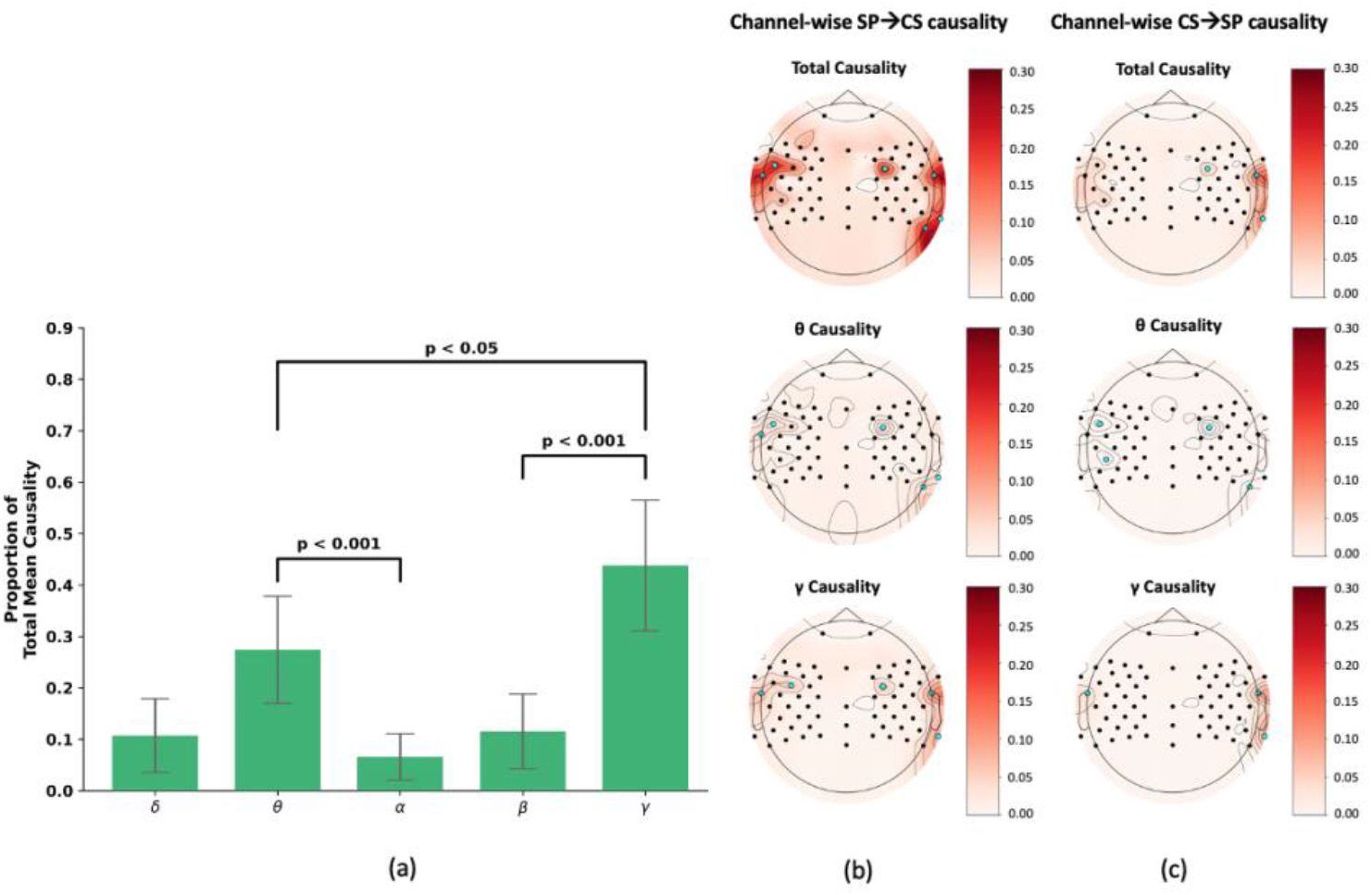
The γ-band produced the greatest spectral causality between SP and CS. (a) Total bi-directional spectral causality in the five neural oscillatory frequency bands represented as the proportion relative to total existing causality. p-values were determined through Wilcoxon rank sum tests. The topographical spectral causality in the SP to CS direction (b) and in the CS to SP direction (c), showing total (top), the θ-band (middle), and the γ-band (bottom) causality. Cyan markings denote significant source channels as determined through z-testing.

Provided that the relationship between SP and CS was predominantly represented by γ activity, we next sought to determine phase-amplitude coupling (PAC) pertaining to this frequency band. Specifically, we examined within-task PAC in SP and CS individually as well as across tasks using SP θ and CS γ. In these analyses, PAC was found to be dynamically modulated in response to word length, with the short word group producing a relatively shorter and weaker PAC and the opposite for the long group (Fig. 10, top row). On the other hand, CS displayed a relatively sparse PAC profile (Fig. 10, middle row), consistent with previous reports (71). Importantly, we found significant SP θ-CS γ cross-task PAC that correlated significantly with that of within-task SP (Fig. 10, bottom row) (Spearman rank, p<0.01).

**Figure 10.**
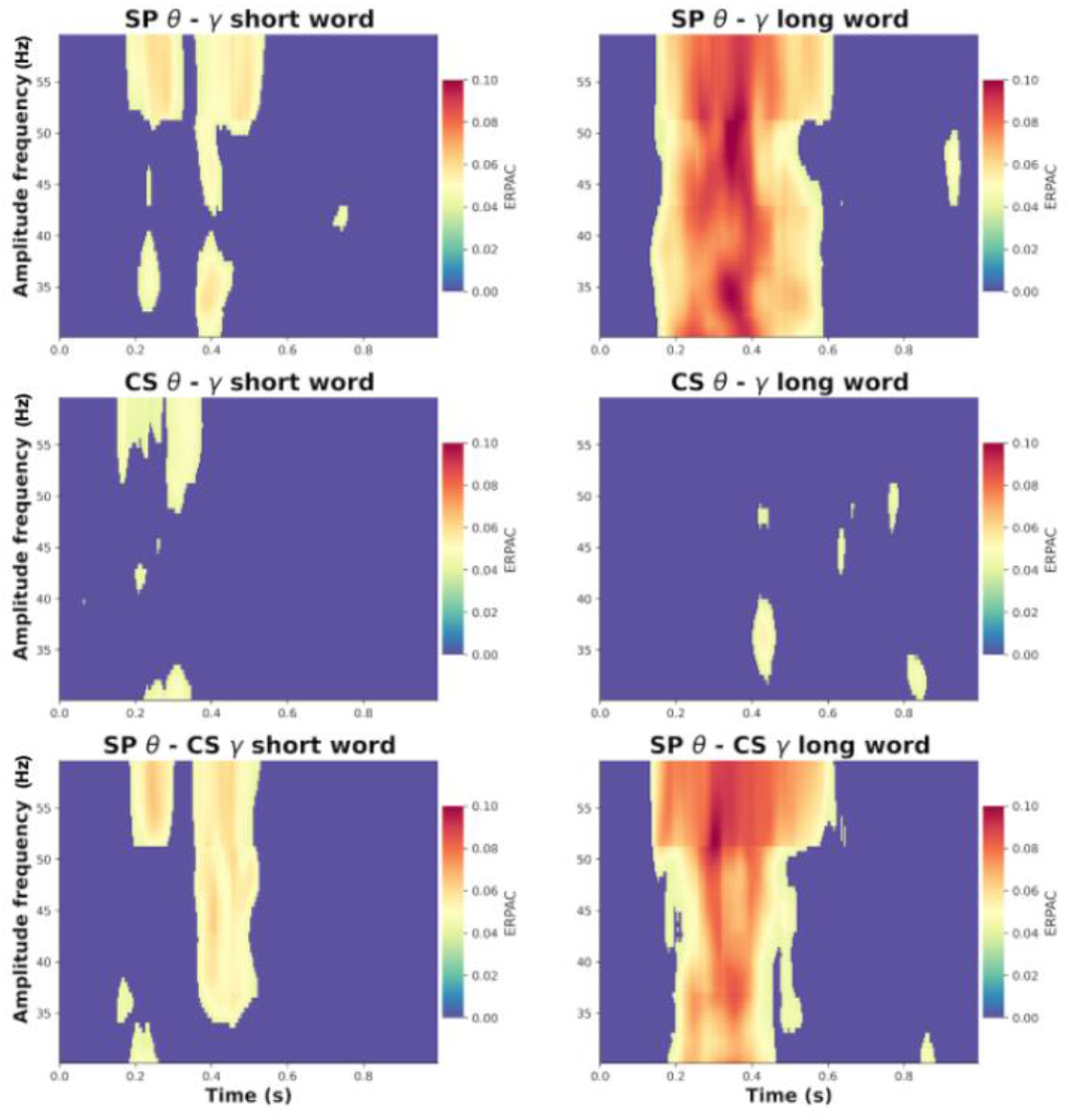
The θ oscillations of SP dynamically couple to both SP and CS γ. The figure depicts PAC between the θ- and γ-bands in each task and word group (short/monosyllabic vs. long/multisyllabic), recorded in the highest common causality-producing channel, TP10. Significant couplings in the time and frequency domains were identified by max-randomization permutation testing. The top and middle rows depict the within-task coupling in SP and CS, respectively. The bottom row depicts the cross-task coupling between SP θ and CS γ, which evidently correlated to that of within-task SP coupling (Spearman rank p<0.01).

Next, we sought to understand how the observed PAC manifests topographically. For this analysis we calculated the PAC within each channel and subsequently averaged across participants. Max randomization Monte Carlo simulations were again conducted. Short and long word SP displayed significant PAC activations in frontal, central/centro-parietal, and temporal channels (Fig. 11 – top row), whereas CS lacked such coupling (Fig. 11 – middle row). Importantly, cross-task PAC (Fig. 11 – bottom row) displayed a topographical PAC profile correlated to that of within-task SP (Spearman rank; short word, r=0.664, p<0.0001; long word, r=0.815, p<0.0001).

**Figure 11.**
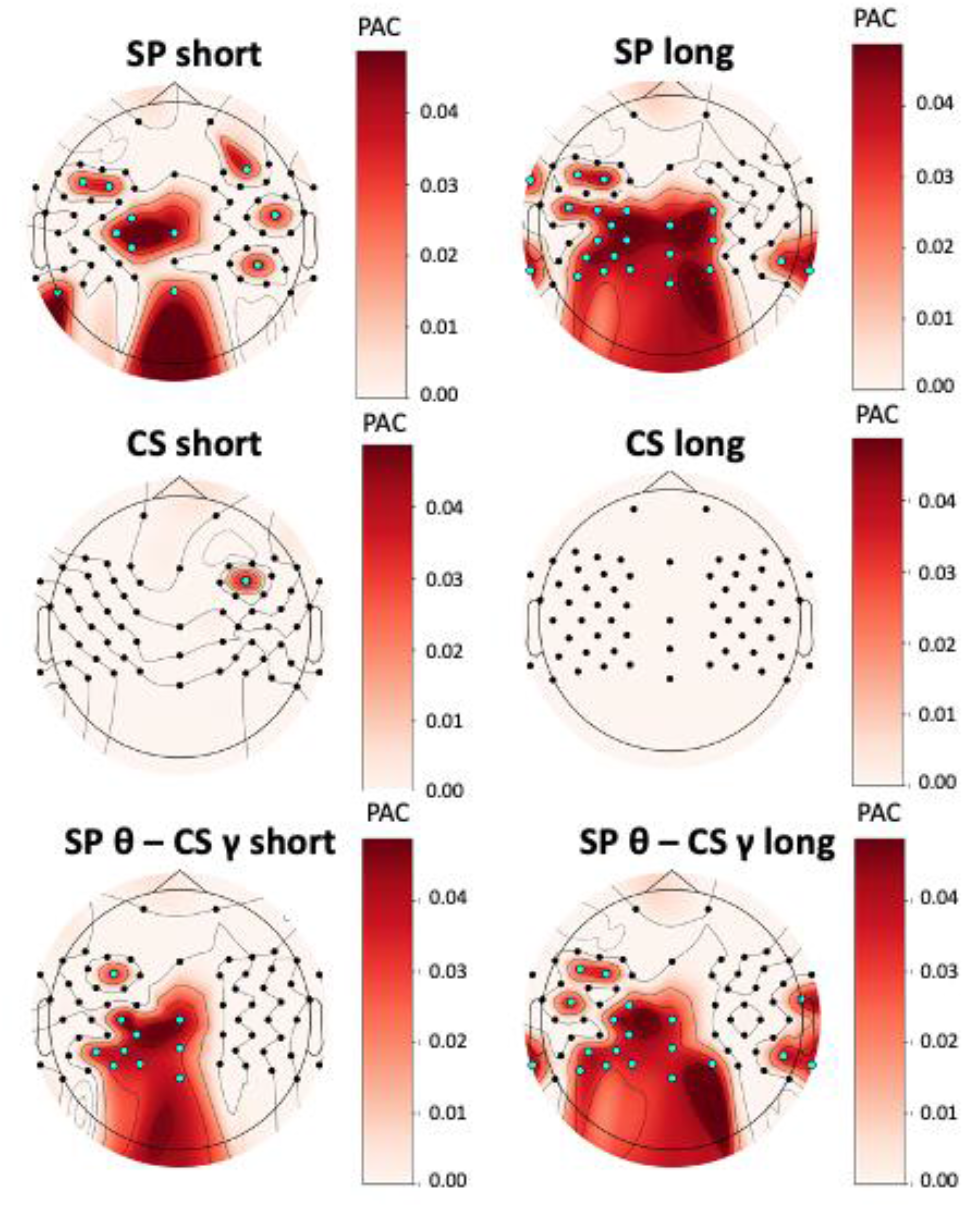
γ activity appears to be spatiotemporally correlated in SP and CS. The figure depicts the PAC topography in SP and CS of short and long word groups, with cyan markings representing significant channels determined through max randomization Monte Carlo simulations.

To assess the relevance of θ and γ oscillations in the current modelling framework, we examined the correlation between accuracy, cross-task PAC strength, γ-band dissimilarity between SP and CS (DTW – dynamic time-warping), root mean squared error of the regression models (RMSE), and proportional change in low-frequency (1-8Hz) power resulting from transformation accomplished through a Ridge regression function derived from SP and CS signals. Considering data from all words and participants, all significant correlations were tested with Spearman tests (p<0.001). We detected the following relationships: 1) accuracy was positively correlated to cross-task PAC strength (Fig. 12a); 2) cross-task PAC strength was negatively correlated with γ-band dissimilarity between SP and CS (Fig. 12b); and 3) γ-band dissimilarity was positively correlated with RMSE (Fig. 12d). Figure 12c depicts the enhancement of low-frequency power in the ERP as a result of SP-CS transformation (Wilcoxon rank sum, p<0.05). Finally, Figure 6e depicts a significant positive correlation between cross-task PAC strength and proportional change in low-frequency power of the transformed CS data relative to the original.

**Figure 12.**
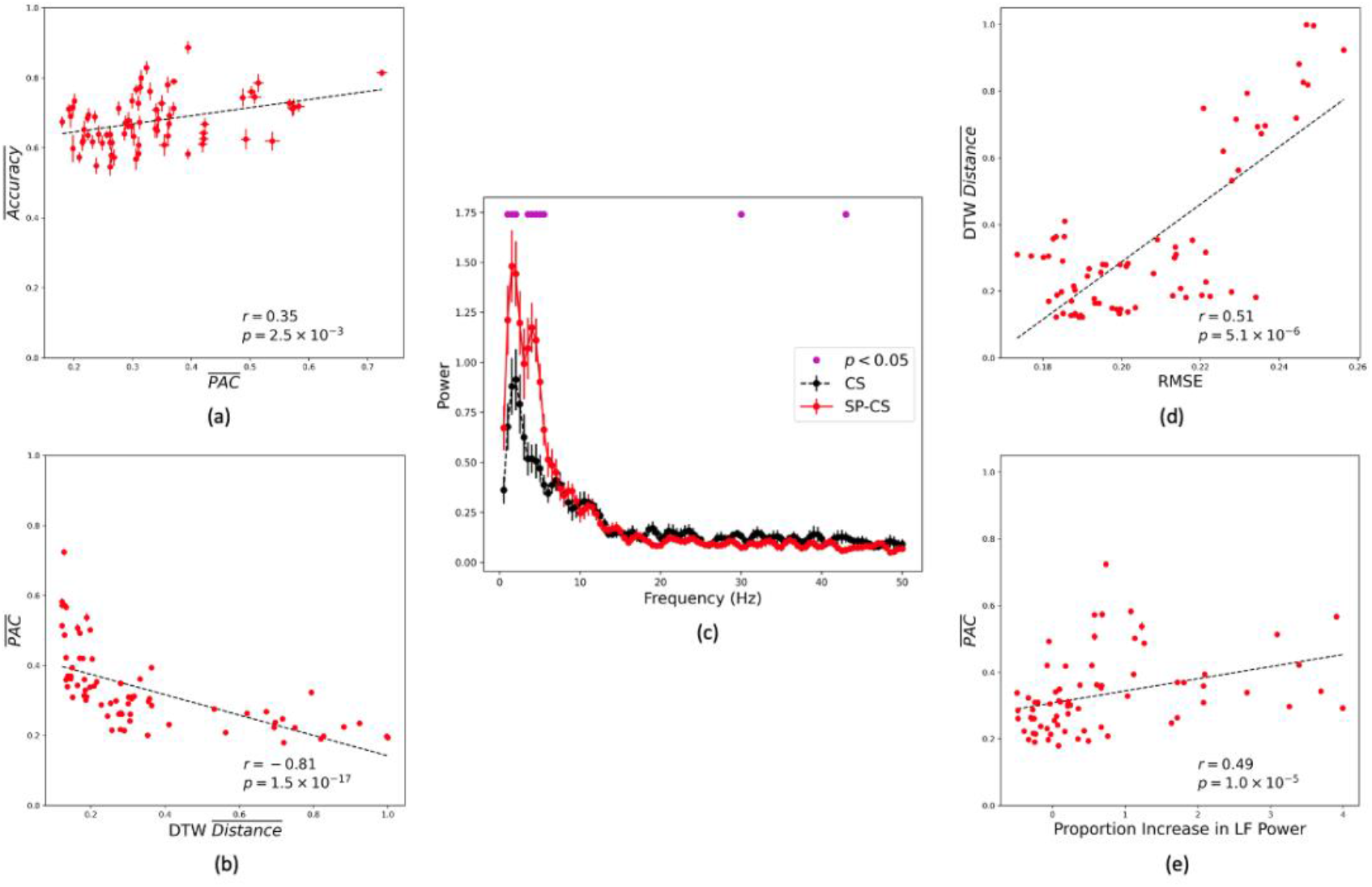
γ-band correspondence may predict model performance and strength of low-frequency power after transformation. Each point represents the average of the depicted quantities for one word for one participant. Dashed lines represent best fit shown for illustrative purposes, with respective p- and r values of Spearman correlation tests. (a) Significant positive correlation between the mean accuracy and mean cross-task (i.e., SP θ – CS γ) PAC strengths. (b) Significant negative correlation between PAC and γ-band dissimilarity (DTW distance). (c) Significant enhancements (indicated by magenta markings) of low-frequency power in the overall (i.e., all words) ERP of transformed CS (i.e., SP-CS) relative to the original data across participants (Wilcoxon rank sum, p<0.05). (d) Positive correlation between γ-band dissimilarity and RMSE estimates of regression (uniform average across channels) (e). Significant positive correlation between cross-task PAC strength and proportional change in low-frequency power of the transformed CS data relative to the original.

## Discussion

In the current study, we designed and demonstrated an online covert speech BCI to classify dyadic trials (sequential SP and CS of the same word). The online BCI linearly transformed signals from CS trials using signals from the preceding SP counterparts through ridge regression and subsequently extracted features using Riemannian geometry. The core finding of the study was that this cross-task modeling approach enabled online discrimination between two words and rest, with ternary online classification accuracies reaching up to 93% and averaging 75.3% ± 9.02%, exceeding the determined chance level (60%). The proposed framework may provide a passive and accessible training method for CS BCIs.

### Cross-condition modelling of SP and CS facilitates classification

Several observations support cross-condition modeling of SP and CS for the discrimination of covertly rehearsed words. In retrospective post-hoc analysis, pure CS classification accuracies fell well below the chance-level (60%), potentially reflective of the low SNR conditions of CS signals alone. Additionally, permuting the CS labels prior to cross-condition modeling significantly reduced classification accuracy, suggesting that the modeling approach leveraged to some degree, inherent SP-CS relationships (discussed later). Binary monosyllabic vs. multisyllabic word classification produced significantly lower mean accuracies than chance, implying that the distinction between classes was not purely based on word duration/length.

### Inter-session variation

The lower online vs. offline accuracies for Participants 4, 7, and 8 stemmed from a significant drop in block-wise online accuracies, namely, in blocks 2 and 3, possibly due to waning engagement or focus from block 2 onward. These participants achieved Block 1 accuracies comparable to those of other participants, suggesting that the current methodology is generally usable by all participants, over short periods of time. The reduction in offline accuracies from session 1 to 2 (for 7 of 10 participants) may be attributable to initial novelty driving more engaged attempts at the tasks and thus eliciting more distinct neural response patterns.

### Superiority of ‘Archbishop’ and ‘Gran’ in online distinguishability

‘Archbishop’ and ‘Gran’ were found to be the most distinguishable word pair consistently across nine out of ten participants, producing the largest inter-cluster distances. The overlap between the topographies from the channel-by-channel perturbation test (Fig. 7d) and word-based r2-scores (Fig. 8c) may attribute distinguishability to a stronger mapping between stimulus features and neural responses for these 2 words. The observed engagement of temporal and central/centro-parietal channel clusters may be indicative of somatosensory transformation relating to auditory representations (34,72). However, greater sample sizes and source modeling potentially with MEG may be required to effectively support these interpretations. Interestingly, ‘Arch’/‘Archbishop’ and ‘Mar’/‘Marketplace’ were grouped into one cluster while ‘Gran’/‘Grandchildren’ and ‘Kin’/‘Kindergarten’ were grouped into another. These groupings may be due to the shared phonology of ‘ar’ in the first cluster, and ‘n’ in the second and may provide evidence of phonological processing manifesting in EEG signals (73).

### θ and γ oscillations in SP and CS: Relation to online classification

The greatest contribution to causality between SP and CS was found in the γ-band, suggesting that their respective EEG signals are spectrally correlated in this frequency range. This finding is perhaps unsurprising, given that low and high frequency EEG signal power has been found to be anticorrelated and that CS preferentially invokes higher frequencies (71,74). In these studies, the authors speculated that the downregulation of low frequencies facilitates the local neuronal processing by higher frequencies. Interestingly, most of the causality between the two tasks was situated in the bilateral temporal regions, potentially indicative of similar auditory representations between tasks. This finding is consistent with reports of γ activity being implicated in corollary discharge (34,75), a mechanism thought to deliver temporally precise and content-specific sensory predictions of speech (76). Therefore, accurate online classification may have been enabled by the shared auditory representations between SP and CS, as manifested in γ activity. This interpretation is further supported by the spatiotemporally correlated PAC profiles for SP θ– SP γ and SP θ–CS γ, which may be suggestive of a common encoding mechanism between SP and CS tasks. In addition, the significant PAC channel clusters appear to overlap with that of the STRF and perturbation test topographies, suggesting that the γ-band likely played a major role in the modelling approach and online classification.

On the other hand, consistent with previous reports (71), CS produced no significant within-task coupling, potentially suggesting that its θ-band may perform a separate encoding mechanism than in SP (74). Although the exact role of θ in CS is not clear, the fact that CS did not produce any significant coupling offers insight into the limitation of this task for BCI control. Importantly, this lack of coupling in CS along with significant lower accuracies in the pure CS classification (Fig. 7c) suggest that the lower frequencies such as the θ-band may contain the salient information necessary for word discrimination. It may thus be reasoned that the bulk of the transformation between the two tasks occurred in the lower frequencies. Indeed, Figure 12c depicts a significant enhancement of low-frequency power following SP-CS transformation. In turn, one could then speculate that the effectiveness of this low-frequency transformation hinged on the relationship between SP θ and CS γ. The correlation between accuracy and cross-task PAC strength (Fig. 12a) seemed to support this conjecture.

## Conclusion

The present study demonstrated that EEG from CS can be classified online using EEG from a SP-CS dyad. The proposed modelling framework significantly enhanced classifier performance beyond that achievable by CS alone. Our findings suggest that *γ*oscillations were pivotal to online CS discrimination. Specifically, the merit of the proposed model may be related to its amplification of low frequency power of CS signals in a manner dependent on the cross-task correspondence of *γ*oscillations. Our results lend credence to the concept of generalizable SP-CS models achieved by a predictive modelling of the lower frequencies in CS.

## List of Abbreviations

CS: Covert Speech
SP: Speech Perception
BCI: Brain-Computer Interface
PAC: Phase-Amplitude Coupling
ERC: Event-Related Causality
EEG: Electroencephalography
STRF: Spectrotemporal Receptive Field
MDS: Multidimensional Scaling
RMSE: Root Mean Sqaured Error
SNR: Signal-To-Noise Ratio

## Declarations

## Acknowledgments

We thank Feny Pandya and Sarah Holman for their help in data collection and Ka Lun Tam for assisting in developing the protocol.

## Competing Interests

The authors have no competing interests to disclose.

### Availability of Data and Materials

The data generated during the current study are not publicly available due to confidentiality constraints, but are available from the corresponding author on reasonable request, and upon approval by the ethics board of Holland Bloorview Kid’s Rehabilitation Hospital.

### Funding

The project has been funded by the Toronto Rehabilitation Institute.

### Ethics Approval and Consent to Participate

All subjects provided consent on their participation in the study. The study was approved by the ethics boards of University of Toronto and Holland Bloorview Kid’s Rehabilitation Hospital.

### Consent for Publication

The authors and subjects of the study gave consent for publication. However, there are no personally identifiable information disclosed in the article.

### Authors’ Contributions

JM – Study design, algorithm development, data collection, analysis, manuscript preparation TC – advised on the above, assisted in manuscript preparation

## Footnotes

1 http://www.blairarmstrong.net/tools/Union_Subtl_CMUPron/index.html

2 https://github.com/pyRiemann/pyRiemann

