## Supporting Information for "Online decoding of covert speech based on the passive perception of speech"

Tom Chau

**This PDF file includes:**

Supporting text

Figures S1 to S2

**Supporting Information Text**

It is important to note that the chance level derived from permuting CS labels was 60%. Since this value may seem high compared to raw chance (33%), we provide an explanation herein. Due to having only two label types from which to permute, the proportion of SP-CS trial pairs for which CS labels remain unchanged despite permutation, was between 0.40-0.64. In other words, the original and permuted label set were not fully independent, yielding a conservative chance level of 60%. This is explained graphically in Figure S7. The mean accuracies were positively correlated to the number of SP-CS dyads with unchanged CS labels, the correction of which resulted in values close to raw chance. Nevertheless, a more conservative chance level affords a more realistic assessment of BCI performance, especially given that the usability threshold, although arbitrary, is conventionally thought to lie between 70-75% (1).

To confirm that the current measure of PAC was in fact sensitive to the specific γ-band patterns present in the signals (as opposed to correlating with any given pattern), we cumulatively perturbed the amplitude of the γ-band signal, one sample at a time, with Gaussian noise (0 mean and variance equal to that of the unperturbed γ amplitude values). The sensitivity curve in Fig. S8 shows that as expected, PAC scores dropped with increasing numbers of noise-perturbed samples in the γ-band signal.

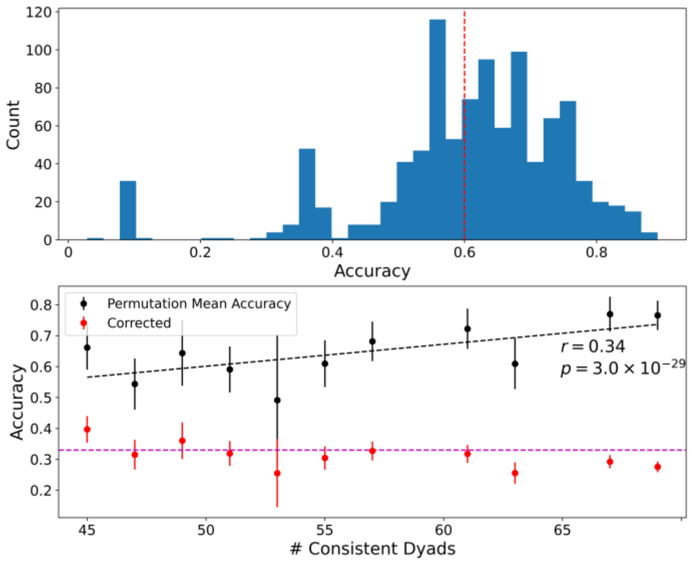


Figure S 1. Classification accuracy after permuting CS labels. The top panel depicts the histogram of accuracies after label permutation, with red dashed line demarcating the mean. The proportion of SP-CS dyads for which CS labels remain unchanged after permutation as 0.4 to 0.64. The bottom panel shows the correlation between the number of mean accuracy and # of SP-CS dyads with unchanged CS labels post-permutation (Pearson, $p<0.05$) in black. In red is the mean accuracy after subtracting from the permutation accuracy the proportion of accuracy due to SP-CS dyads with unchanged labels. The magenta dashed line demarcates raw chance level (33%).

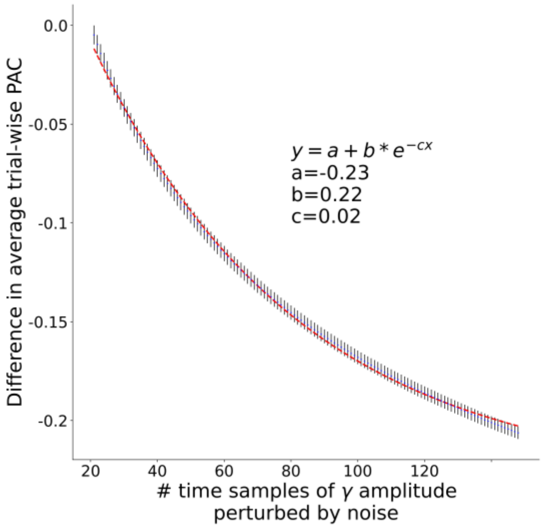


Figure S 2. γ sensitivity curve. PAC values were determined by progressively perturbing the γ-band with random gaussian noise (0 mean, variance equal to unperturbed γ amplitudes) one sample at a time and recording the drop in PAC strength with each subsequent step. PAC strength falls exponentially with increasing number of samples perturbed. Bars indicate standard error.
